# NF-κB Epigenetic Attractor Landscape Drives Breast Cancer Heterogeneity

**DOI:** 10.1101/2024.07.10.602798

**Authors:** Francisco Lopes, Bruno R. B. Pires, Alexandre A. B. Lima, Renata Binato, Eliana Abdelhay

## Abstract

Heterogeneity in breast cancer (BC) subtypes contributes to therapy resistance and recurrence. Subtype heterogeneity arises from stochastic genetic and epigenetic changes, phenotypic plasticity, and microenvironment-driven selection during tumor evolution. Here, we investigate how NF-κB epigenetic variability contributes to HER2^+^ BC progression. Using RNA-seq, we quantified NF-κB, TWIST1, SIP1, and SLUG expression in two BC cell lines: HCC-1954 (HER2^+^) and MDA-MB-231 (TNBC). Next, we built and calibrated a gene regulatory network model reproducing transcriptional interactions among these genes. The model’s epigenetic landscape displays two attractor basins that reproduce the HER2^+^ and TNBC expression profiles. Validation was performed using DHMEQ-treated cells, published patient and in vitro data. Stochastic fluctuations in NF-κB levels induce spontaneous, irreversible transitions from HER2^+^ to TNBC states at variable times, contributing to heterogeneity. These transitions are mediated by an unstable intermediate state that provides a noise-sensitive route. Mutations or drugs altering NF-κB availability reshape basin sizes, altering basin sizes and transition probabilities. Our work refines the attractor landscape framework, linking NF-κB dynamics to BC heterogeneity, supporting more accurate classification, prognosis, and treatment strategies.

## INTRODUCTION

To understand how cell differentiation occurs rapidly during embryonic development, Waddington proposed his famous metaphor, which consists of the sequential events of cell differentiation compared to a ball rolling down a bifurcated valley (1), Fig. 1A. Later, using the dynamic systems theory, Kauffman (2,3) proposed the concept of cell types as attractors. According to this concept, all cell types, including cancer cells, correspond to different attractors in a multidimensional space. Transitions between different attractors are thought to be induced by biochemical stochastic fluctuations, with gene mutations potentially facilitating these transitions to cancer cells by lowering the barriers separating the attractors. Building on this foundation, Wang and collaborators (4) have recently proposed a systematic approach to determine a probability landscape, leading to a quantitative characterization of Waddington’s epigenetic landscape, thus transforming it from a metaphor into a more quantifiable framework. Bhattacharya and collaborators (5) proposed an alternative, simpler approach, where a quasi-potential surface is directly derived from the network ODE system. Additionally, computational strategies such as SCUBA (single-cell clustering using bifurcation analysis) have described lineage relationships between cells at various developmental stages by identifying bifurcation events that lead to the emergence of new attractor states during differentiation (6). However, despite the conceptual elegance of Waddington’s landscape and Kauffman’s attractor model, they often lack precise molecular mechanisms underlying these processes and fail to fully explain the nature and timing of the fluctuations that induce transitions. More detailed molecular models are needed to bridge the gap between abstract landscapes and actual biological pathways.

**Figure 1.**
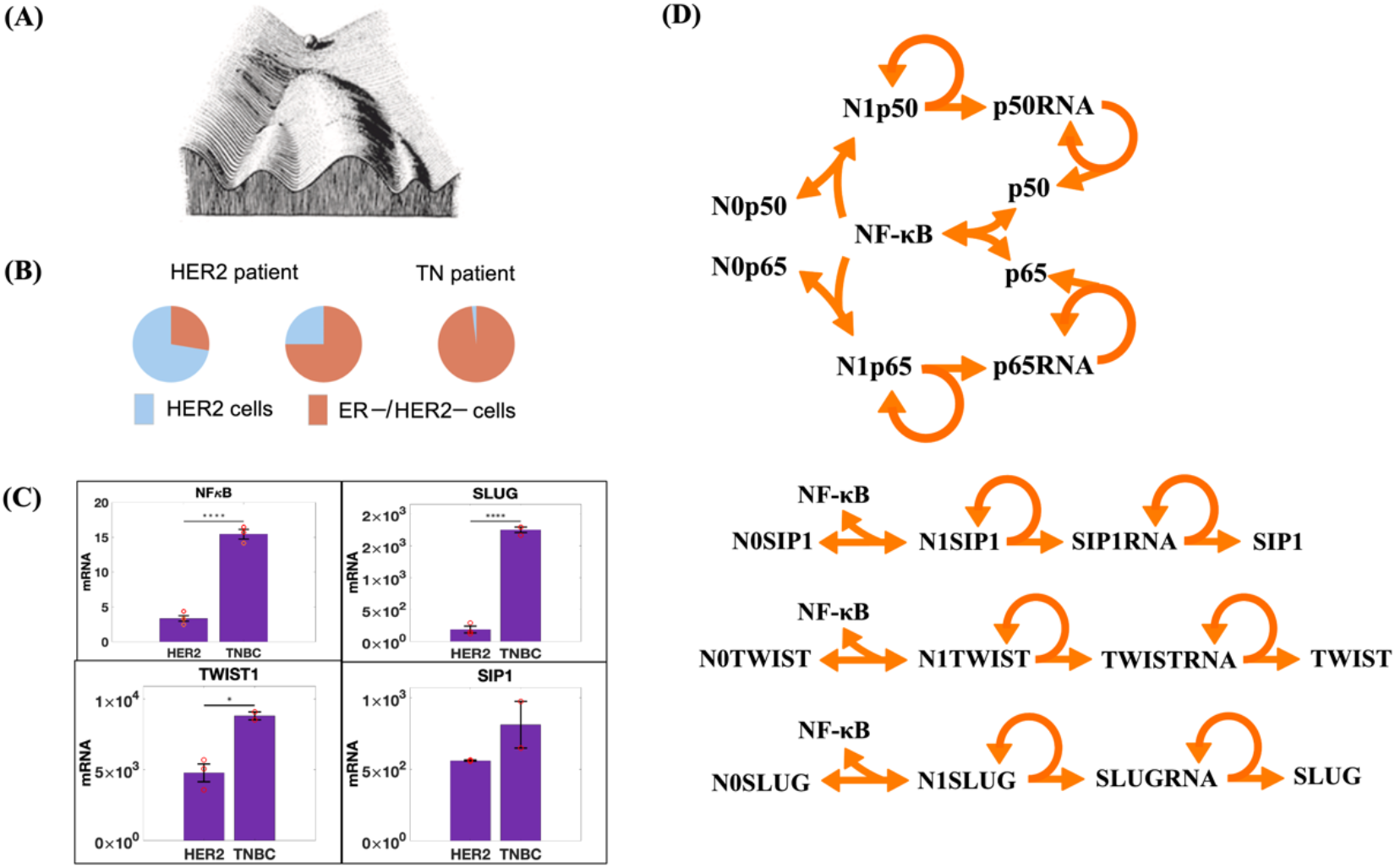
Model for the activation of SLUG, SIP1, TWIST1, and NF-κB (subunits p50 and p65). (**A**) Waddington epigenetic landscape metaphor. (**B**) BC heterogeneity determined by scRNAseq (6): proportion of HER2^+^ and ER^-^/HER2^-^ cells in two HER2 and one TN classified patient. (**C**) Quantitative PCR (qPCR) analysis of NF-κB, TWIST1, SLUG, and SIP1 expression levels in HCC-1954 (HER2^+^) and MDA-MB-231 (TNBC) cell lines, ****, ***and * indicates e^-4^, e^-3^ and 2e^-1^ statistical significance, respectively. RNA levels were estimated from our qPCR data compared to the estimated number of RNA molecules in a mammalian cell (Table S1). (**D**) Representation of model reactions: N0X or N1X indicates whether the regulatory region of a given gene X is empty or occupied by a NF-κB dimer, respectively. Gene X can be SLUG, SIP1, TWIST1, p50 or p65. Gene activation is described by a three-step reaction: a reversible reaction for NF-κB binding to the gene regulatory region (NFκB + N0X ↔ N1X), and two irreversible reactions for RNA (N1X → N1X + XRNA) and protein synthesis (XRNA → XRNA + X). Constitutive synthesis and degradation of RNAs, and protein degradation are also assumed. Detailed reactions on Fig. S1.

BC is commonly classified into three subtypes: luminal, characterized by high expression of estrogen receptors (ER) and often progesterone receptors (PR); HER2-positive (HER2^+^), marked by overexpression of the epidermal growth factor receptor 2 (HER2); and triple-negative breast cancer (TNBC), which lacks expression of ER, PR, and HER2 (7). Using single-cell RNAseq (scRNAseq) to classify cells from the same BC patient sample, Chung and collaborators (8) found remarkable heterogeneity (Fig. 1B). Wu et al (9) also revealed significant heterogeneity in samples from luminal, HER2^+^ and basal (TNBC) patients using scRNAseq data as well.

The NF-κB family consists of five subunits (RelA/p65, NF-κB1/p50, NF-κB2/p52, c-Rel, and RelB) that form various homo- and heterodimers, with p65/p50 being the most prevalent (10). In resting cells, NF-κB is bound and inhibited by IκB complexes, consisting of IκBα, IκBβ, and IκBε subunits. The activation of NF-κB signaling has been associated with numerous pathological conditions, including chronic inflammation (11), heart (12) and immune (13) diseases, and various types of cancer (14-19). In cancer cells, NF-κB signaling is involved in diverse mechanisms like cancer initiation, uncontrolled proliferation, metastasis, and therapy resistance (19). At the transcriptional level, we have previously shown that NF-κB directly activates epithelial-mesenchymal transition (EMT) transcription factors SNAI2/SLUG, ZEB2/SIP1, and TWIST1, playing a critical role in EMT, which is a characteristic of HER2^+^ and TNBC cells (20).

To investigate NF-κB role in BC progression, we built a gene regulatory network (GRN) model depicting NF-κB regulation of itself and the target genes TWIST1, SLUG, and SNAIL (20-22). The model shows that the cell attractor landscape has two stable states, each exhibiting the gene expression levels we determined for HER2^+^ and TNBC breast cancer subtypes. Stochastic simulations induce spontaneous transitions from HER2^+^ to TNBC basins. No transition from TNBC to HER2^+^ was verified in any cell simulation, indicating a preferential direction of the transitions, which is compatible with the TNBC subtype being a more advanced cancer stage than HER2^+^. This preferential direction is related to the size of both basins in the cell attractor landscape. Interestingly, the correlation between the p65 and p50 RNA in TNBC is lower than in HER2^+^, in accordance with Chung et al (8). This is related to the more symmetric shape of the TNBC epigenetic basin compared to that of the HER2^+^ subtype. It suggests that using correlation to describe gene interactions might be less effective in more severe subtypes. Single-cell simulations show that even identical cells exhibit the HER2^+^ to TNBC transition at different times. This indicates that stochasticity works at both protein/RNA concentrations and the tissue identity, playing a critical role in cell type heterogeneity in a tumor site.

## RESULTS

### A GRN reproduces NF-κB dynamics in HER2+ and TNBC cells

To investigate the molecular mechanism underlying NF-κB role in BC progression, we performed RT-qPCR assays to characterize NF-κB expression levels, along with its EMT-related target genes TWIST1, SNAIL, and SLUG, in two well-established breast cancer cell lines representing the HER2+ and TNBC subtypes: HCC-1954 and MDA-MB-231, respectively (**Fig. 1C**). We then built a GRN model (**Fig. 1D, Fig. S1**; *Materials and Methods*) in which the NF-κB heterodimer (p65:p50) activates the expression of TWIST1, SNAIL, SLUG (20) as well as both NF-κB subunits p50 and p65, whose positive feedback has already been described in the literature (21,22). According to chemical reaction network theory (23,24), the model can exhibit two stable stationary states. To evaluate the model ability to replicate gene expression levels in both BC subtypes, we calibrated it using the RNA levels of NF-κB, TWIST1, SNAIL, and SLUG across both subtypes simultaneously. Due to the model bistable behavior, it successfully reproduces the expression levels of both subtypes with a single parameter set, where each subtype corresponds to a distinct stationary state (**Fig. 2A**).

**Figure 2.**
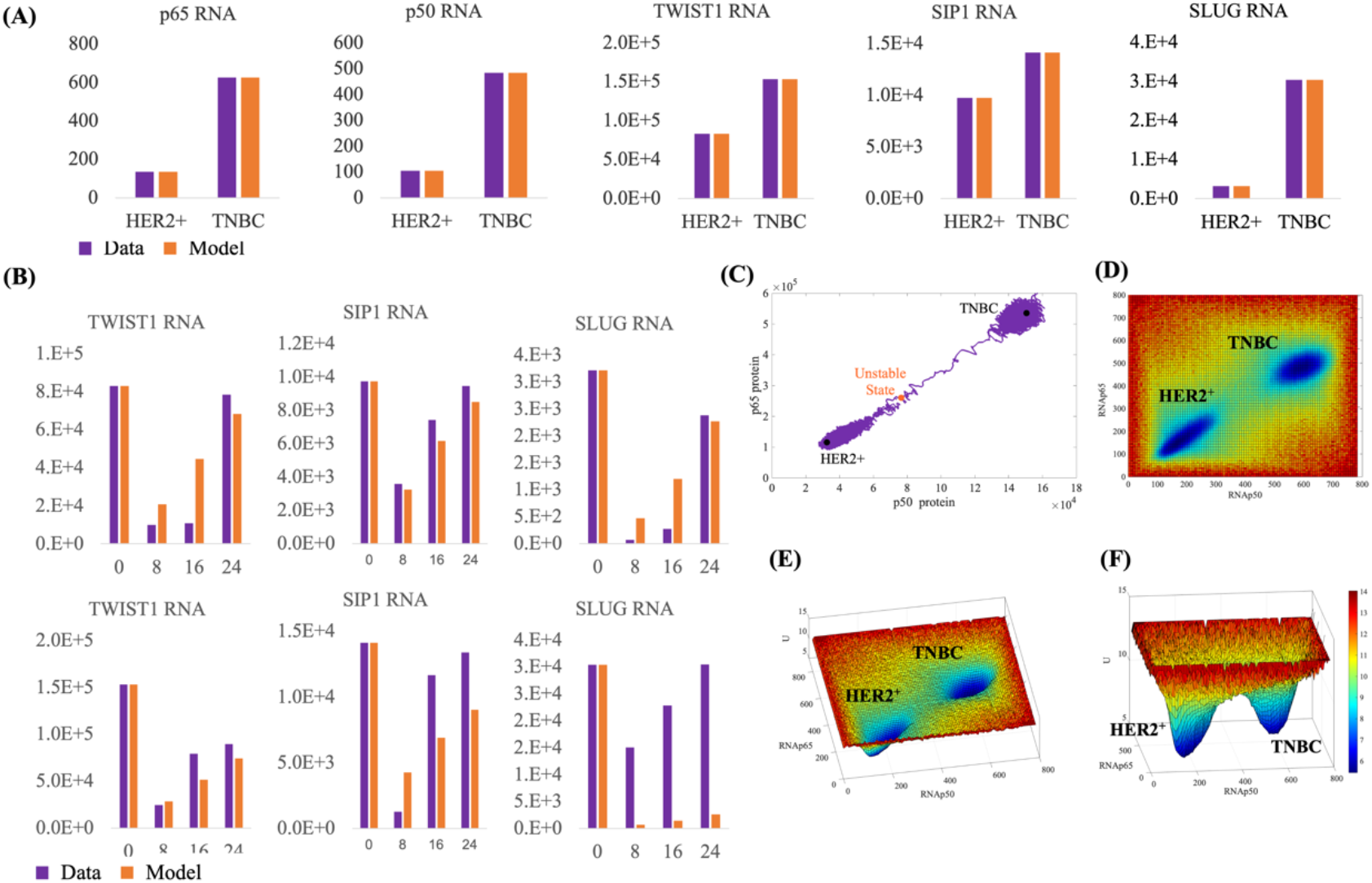
The epigenetic attractor landscape shows an attractor associated with each BC subtype. **(A)** Model calibration to the estimated RNA molecule numbers of NF-κB subunits (p50, p65) and transcription factors (SLUG, SIP1, and TWIST). Orange bars represent the same data as shown in Fig. 1C, while purple bars depict the calibration of the GRN model. (**B**) Fine-tuning of model parameters to replicate the temporal dynamics of expression level recovery in HER2+ (upper panel) and TNBC (lower panel) cell subtypes after treatment with the NF-κB inhibitor DHMEQ. (**C**) The three stationary states in the p50-p65 phase space, with the purple line showing the trajectory of a single-cell simulation undergoing the HER2+ to TNBC transition. (**D-F**) Three different perspectives of the model attractor basins.

To assess the model ability to replicate the temporal dynamics of these genes, we measured the recovery time of their expression levels in both HER2^+^ and TNBC subtypes following treatment with the NF-κB inhibitor DHMEQ (**Fig. 2B**). The model parameters were then fine-tuned to reproduce this RNA recovery without compromising the calibration shown in **Fig. 2A**. The model successfully replicated recovery times comparable to those observed in both cell lines (**Fig. 2B**). The only exception is SLUG, where experimental recovery times differ between the HER2+ and TNBC subtypes. This suggests that additional genes may contribute to the faster SLUG recovery observed in TNBC. Notably, the experimentally determined high SLUG levels at 8 hours in TNBC cells (purple bars) compared to HER2^+^ cells indicate the presence of other transcriptional regulators beyond NF-κB in TNBC. Since these factors are not included in the GRN model, this effect could not be reproduced.

### The BC GRN exhibits an attractor landscape with two basins

To describe the model phase space, we solved the algebraic equations derived from the Ordinary Differential Equations (ODEs, **Fig. S2**) finding three stationary states (**Fig. 2C**). Both black circle states have seven eigenvectors with negative eigenvalues and two with null eigenvalues. Between these stable states, there is one unstable one. We then built the epigenetic attractor landscape by performing 10 thousand stochastic simulations, starting at different uniformly-distributed positions in the p50*-*p65 RNA phase space (**Fig. 2D-F**). This surface exhibits two basins of attraction, each containing the stationary position of one of the BC cell subtypes. The TNBC basin is more symmetric than the HER2^+^ one, as shown in Fig. 2C. Also, the unstable position is closer to the HER2^+^ basins than to TNBC one. All these characteristics are discussed in depth in the following sections.

### Stochastic Fluctuations induce the HER2+ to TNBC Transition

According to the Kauffman’s cancer attractors model (2,3), stochastic fluctuations in gene expression drive the transition between attractor states, which are associated with different cell types. To test if our modeling corroborates Kauffman’s hypothesis, we plotted the trajectory of a 38-year single-cell stochastic simulation in the p50*-*p65 protein phase space (**Fig. 2C**). The simulation follows a trajectory that visits different positions around the equilibrium state in the HER2^+^ attractor (**Fig. 2C, Movie S1**). These positions have a strong asymmetrical distribution, resembling the shape of the HER2^+^ basin and suggesting confinement within it (**Fig. 2D**). While confined to this basin, the trajectory is restricted to the region between the HER2^+^ stable and the central unstable state; however, if the trajectory passes through the position of the unstable state, it is strongly attracted towards the eventually TNBC basin. Once there, the trajectory becomes more symmetrical than in the HER2^+^ basin, resembling the shape of the TNBC basin. We see that the stochastic trajectory is spontaneously moving from the HER2^+^ to the TNBC basis of attraction. This result corroborates the Kauffman’s cancer attractors hypothesis showing that, at the microscopic level, stochasticity in the concentrations of proteins and RNAs induces the transition between the attractor basins, which are associated with the HER2^+^ and TNBC subtypes.

Given that the asymmetrical shape of the trajectory in both basins can be crucial for the correlation between positions in the phase space, we decided to compare the correlation between p50 and p65 RNA levels from our simulations with a cohort of basal and HER2^+^ non-treated Asian patients (n=51, (25, 26). In Fig. 3A and B we show a set of simulations exhibiting Spearman correlation (Sc) of 0.66 (*p* = 3.7e^-3^) for HER2^+^ and 0.56 (*p* = 5.8e^-4^) for TNBC simulations, respectively. These results align with the data from non-treat patients (23), that show Sc = 0.60 (*p* = 1.0e^-2^) for HER2^+^ and Sc = 0.56 (*p* = 6.5e^-4^) for TNBC (Fig. 3). They also align with Chung et al (8), who found higher cell-to-cell correlation in TNBC than in HER2^+^.

**Figure 3.**
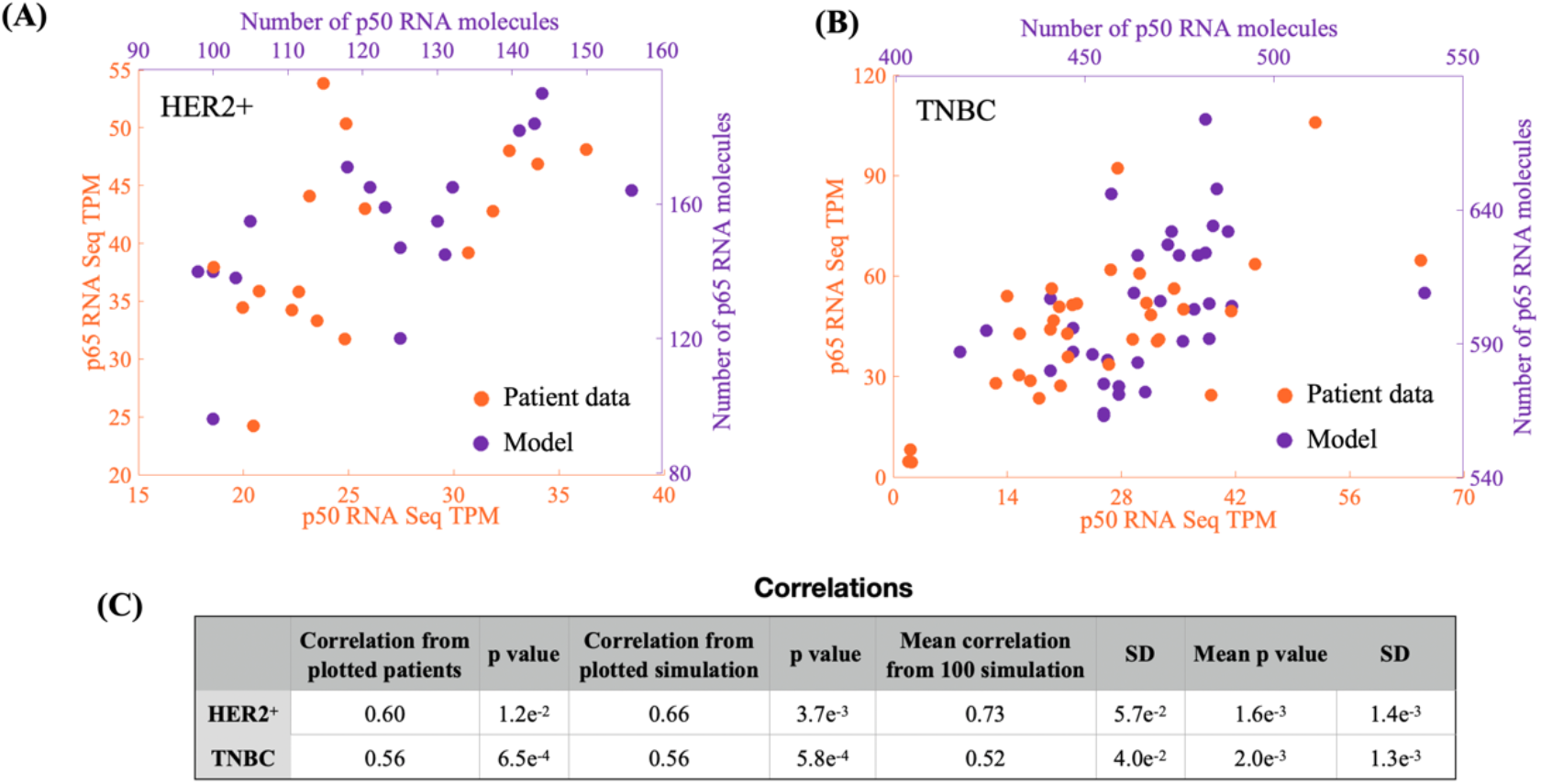
Validation of GRN with data from untreated patient cohort. (23, 24). Orange represents data, and purple represents model simulations. (**A**) Comparison of data from 17 HER2^+^ patients with 17 randomly selected single-cell stochastic simulations from a set of 120 simulations. **(B)** Comparison of data from 34 TNBC patients with 34 single-cell simulations, similar to (A). (**C**) Correlation measures for data in (A) and (B), alongside mean correlations from 100 randomly selected simulations (see *Materials and Methods*).

### The unstable state mediates the HER2+ to TNBC transition

To understand what drives the cell dynamics to escape from the HER2^+^ toward the TNBC basis of attraction, we projected all seven eigenvectors with non-zero eigenvalues into the 3D p50*-* p65*-*NFκB phase space, Fig. 4A (upper panel) and Fig. S3. These weaker eigenvectors are more susceptible to be overcome by stochastic fluctuations. This susceptibility is key in understanding the transition trajectory depicted in Fig. 2C.

**Figure 4.**
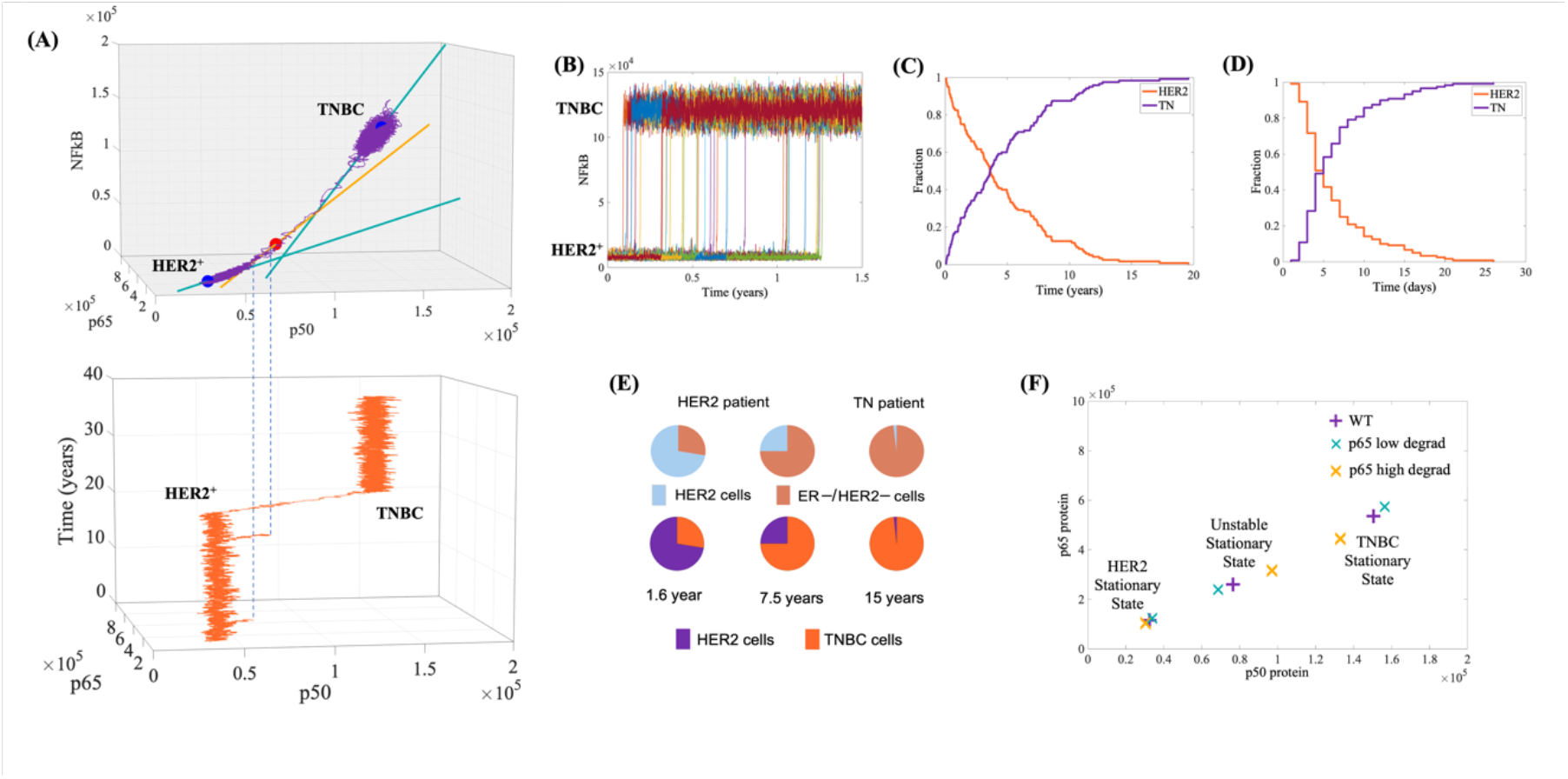
The unstable state facilitates the transition from HER2+ to the TNBC subtype. (**A**) Upper panel: Projection of the weak eigenvectors within the p65-p50-NFκB phase-space. The purple line depicts a single-cell simulation trajectory. Lower panel: Temporal variation of p50 and p65 protein levels in the same simulation. (**B**) Spontaneous transitions from HER2+ to TNBC occurring at various time points over a 1.5-year period, collected from 120 single-cell stochastic simulations. These simulations show a 100% transition rate over 20 years (as shown in **C**). (**D**) Reducing p65 degradation by 2.5% significantly decreases the time required to reach 100% transitions to just 26 days. (**E**) Snapshots from different time points across the 120 simulations (shown in C) recapitulate the tumor heterogeneity reported by Chung et al. (6). (**F**) p50-p65 phase-space displaying the model’s stable positions under conditions of reduced or increased p65 degradation (see Table 1 for details on the shifting measures).

Initially, the trajectory is restricted to the HER2^+^ attractor, where it started (Fig. 4A, Movie S2). It then fluctuates along the green eigenvector (weak, eigenvalue = - 6.0e^-3^) associated with the HER2^+^ stationary state. The lower panel in Fig. 4A shows a 3D plot of the time course from the same simulation. The dotted blue lines between both panels indicate that while the trajectory mainly fluctuates near the HER2^+^ stable position, it moves toward the orange (weak) eigenvector twice but falls short of reaching the unstable position (red sphere). Only on the third instance does the trajectory reach the unstable position, moving past it and into the TNBC basin of attraction.

**Table 1.**
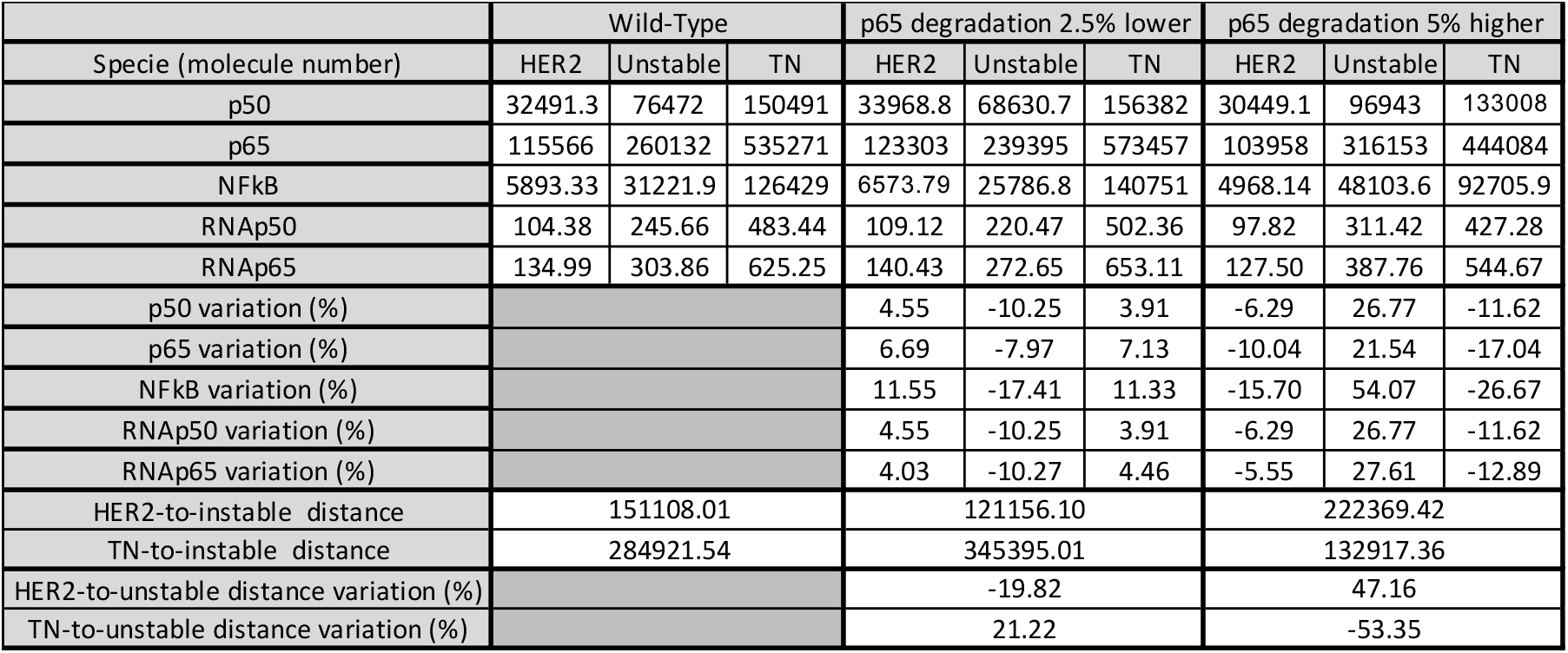
Effects of mutations on the size of attractor basins. A comparison between Wild-Type and mutant stationary states. The table presents the percentage variation in stationary concentrations of NF-κB and its subunits p50 and p65 due to changes in the p65 degradation rate. Additionally, it details the changes in distances from the HER2+ and TNBC stationary states to the unstable states.

This dynamic effectively utilizes the weak eigenvectors susceptibility to fluctuations. Although the red eigenvector (eigenvalue = + 2.5e^-3^) tends to drive the trajectory away from the unstable red position, the fluctuations eventually prevail. Once past the unstable point, the trajectory follows the direction of the orange eigenvector, moving away from the red circle and towards the TNBC stable position. When the trajectory shifts from the orange to the green eigenvector (eigenvalue = - 3.2e^-3^), it continues toward the TNBC position. Once there, it fluctuates around the TNBC stable position. Notably, none of an additional set of 120 simulations over 20 years, starting in the TNBC stable position, showed a transition back to the HER2^+^ basin.

### Spontaneous transitions from HER2+ to TNBC generate tumor heterogeneity

To explore whether microscopic stochasticity contributes to macroscopic heterogeneity in cell collectives (tissues), we conducted 120 identical single-cell simulations over a 20-year model period, each starting from the central stationary state of the HER2^+^ subtype. Assuming no cell-to-cell signaling, all HER2^+^ cells eventually transitioned to the TNBC subtype (Fig. 4B, C), with each cell transitioning at random times and without reverting back. These results suggest that, within a tissue, stochastic fluctuations in gene expression can drive significant epigenetic variability among genetically identical cells. This epigenetic variability, caused by asynchronous transitions from the HER2^+^ to the TNBC subtype, results in the coexistence of different cell subtypes within the same tissue, a key feature of intratumoral heterogeneity.

To validate our results, we compared our 120 stochastic simulations with Chung et al’s study (8), which involved scRNAseq of 515 cells from 10 chemotherapy-naive BC patients and revealed significant intratumoral heterogeneity. In their classification of individual cells, they reported cases such as a sample from a HER2^+^ patient with approximately 16% or 45% of cells categorized as TNBC, and a TNBC patient with 1% of cells classified as HER2^+^ (Fig. 4E, upper panel). By compiling the number of cells that exhibited a transition from HER2^+^ to TNBC at 1.6, 7.15, and 15 years in the model timeframe, we successfully reproduced the relative proportions of cells in each subtype as observed in their study (Fig. 4E, lower panel).

### Mutations increase transition probabilities by changing basin sizes

Kauffman and collaborators (2, 3) have previously proposed that mutations lowering the epigenetic barrier facilitate transitions between adjacent attractors. Recently, Ren et al. (27) demonstrated that changing FBXW2, a protein affecting p65 degradation, alters cell proliferation and drug resistance: reducing FBXW2 increases p65 and cell growth, while increasing FBXW2 decreases p65 and inhibits growth, also affecting paclitaxel resistance. In our study, we investigate the impact of these FBXW2 mutations by simulating changes in the degradation rate of the p65 protein. In HER2^+^ cells, decreasing p65 degradation by 2.5% raised its levels by 6.7%, while increasing degradation by 5.0% reduced p65 levels by 10.0% (Fig. 4F, Table 1). These changes in p65 degradation altered the HER2^+^ attractor basin size: increasing p65 degradation reduced the distance between the stable and intermediate unstable state by 19.82%, while decreasing it increased this distance by 47.16% (Fig. 4F, Table1). Concurrently, the TNBC attractor basin size was inversely impacted by p65 degradation changes: a 2.5% decrease in p65 degradation increased the distance to the TNBC state by 21.22%, while a 5.0% increase reduced this distance by 53.35%.

The above-described effects in the size of the attractor basin suggest explanations for the effect of varying p65 degradation as reported by Ren et al. (27). Specifically, the reduction in p65 degradation shrinks the HER2^+^ attractor basin, facilitating the transition to the state of higher p65 levels, increasing cell proliferation and contributing to tumor progression. Conversely, the model predicts that increased p65 degradation would expand this basin, reducing the likelihood of this transition and thereby decreasing cell proliferation.

To verify whether the above-described effects on the basin sizes influence transition probabilities, we carried out 120 stochastic simulations with p65 degradation reduced by 2.50%. All cells moved to a higher p65 level basin within 30 days, aligning with Ren et al. (27) findings on enhanced cell proliferation; Reinforcing that mutations reducing the size of an attractor basin increase the probability of a cell departing from this attractor. In contrast, another set of 120 simulations with a 65.0% increase in p65 degradation showed no transitions over 20 years. On the other hand, a different series of 120 stochastic simulations with p65 degradation increased by 5.0%, shows no transition even over a span of 20 years. It indicates that mutations that increases the size of an attractor basin decreases the probability of a cell to departure from this attractor. In this case, reducing cell proliferation, in agreement with Ren et al. (27). Taken together, these findings highlight the potential of p65 as a target for Targeted Protein Degradation, an emerging concept in drug discovery (28).

## CONCLUSIONS

We propose a model that is sufficiently small to allow for a detailed analysis based on dynamic systems theory, yet large enough to encompass the key biological mechanisms behind the gene expression levels it intends to describe. In addition, we performed the model calibration in two independent steps. First, we calibrated the model against cell data at the stationary states, followed by a time-dependent calibration to reproduce the dynamics of the gene expression recovery levels. Interestingly, the model could not reproduce the dynamic behavior with the same accuracy as it did for the static states, indicating that the effect of additional regulators was missing. This outcome suggests that our model was not constructed with an unnecessary number of parameters, which could have resulted in overfitting.

Remarkably, such a small model could offer explanations not only for the *in vitro* data it was calibrated against but also for other *in vitro* experiments (27) and patient-derived data (8, 9, 25). This includes explaining the p65-p50 RNA correlation in HER2^+^ and TNBC, in both bulk and single-cell data. In fact, our modeling strategy mirrors the approach of a biologist who selects specific mechanisms to replicate in an *in vitro* assay, aiming to understand the broader *in vivo*, systemic behavior.

By showing how stochasticity at the molecular level leads to heterogeneity in cells and tissues, our work provides an explanation for recent findings about BC heterogeneity (8, 9). This result can potentially impact clinical research by contributing to BC classification, which plays a critical role in determining patient-specific treatment. This can also contribute toward personalized medicine.

By simulating variations in NF-κB dimer availability, our model serves as a platform for preliminary drug testing, particularly for strategies based on Targeted Protein Degradation. Additionally, the current version of the model is ready for diverse expansions. For instance, including target genes such as TWIST, SIP1, and SLUG can enhance our understanding of their roles in EMT, contributing to BC metastasis.

The finding that stochastic fluctuations can induce cell-subtype transitions is particularly interesting. It not only proposes an explanation for tumor heterogeneity but also supports the epigenetic attractor landscape model proposed by Waddington and Kauffman, specifically the association between attractor basins and cellular subtypes. Moreover, our model introduces two enhancements to these well-established general models. First, the model proposes that the key factor influencing how mutations affect transition probability lies in the size of the attractor basins. This offers a valuable tool for understanding and predicting the conditions necessary to trigger or prevent these transitions, because the size of the basins can be estimated from the relative distance between the stable states. Second, the model provides a detailed description of how eigenvectors contribute to transitions between basins of attraction. It proposes a specific pathway for transitions between the subtypes of these basins and offers an explanation for their irreversible nature. This irreversibility arises from two key factors: the relative weakness of the eigenvectors positioned between the stable states and the distance between these states. The weakness of the eigenvectors allows fluctuations to override their directionality, while the distance between the states affects the likelihood of a fluctuation being strong enough to push the system across the unstable position and into another attractor basin. These insights not only enhance our understanding of tumor dynamics but also highlight potential therapeutic approaches. Developing strategies to prevent or slow these stochastic transitions could offer a way to reduce intratumoral heterogeneity and improve treatment outcomes.

Finally, this work highlights the advantages of an open-box model-building strategy. This approach utilizes relatively small models, which are well-characterized by dynamic systems theory, to accurately reproduce specific cellular processes. These smaller models can then be interconnected to create larger, more comprehensive models that capture the intricate interactions between various cellular processes. Successful examples of this strategy are already evident in the literature (29-32), including its effective integration with successful black-box approaches (33).

## MATERIALS AND METHODS

### Cell culture and Real-Time Reverse Transcription Polymerase Chain Reaction (RT-qPCR)

The human BC cell lines MDA-MB-231 and HCC-1954 were cultured as previously described (20). Total RNA was isolated using TRIzol reagent (Thermo Fisher) according to the manufacturer’s instructions. Then, 2μg of RNA were treated with the DNase Amplification Grade I Kit (Thermo Fisher) and reverse transcribed into cDNA using the Superscript-II kit (Thermo Fisher) following the manufacturer’s protocol. RT-qPCR was performed with SYBR Green Master Mix (Bio-Rad) in a Rotor-Gene Q (Qiagen) under previously reported conditions (20). Each sample was examined in triplicate. ACTB and GAPDH were used as the reference genes for the RNA levels. Fold-expression was calculated according to the ΔΔCt method (34).

### Building the Gene Regulatory Network

In our model, gene activation is described by a set of three-step reactions (35): a reversible reaction where the transcription factor (TF) binds to the gene regulatory region, followed by two irreversible reactions for RNA and protein synthesis (Fig. 1D and Fig. S1). This strategy is consistently applied to describe the transcriptional activation of all five genes: p65, p50, TWIST1, SLUG, and SIP1. The model is completed by generic source and decay reaction for all proteins and RNAs. We applied the law of mass action to derive a set of ODEs that describe the model dynamics (Fig. S2). The cell RNA levels were estimated from our RT-qPCR data (Fig. 1C) compared to the estimated number of RNA molecules in a mammalian cell (Table S1). To ensure model accuracy, we assumed that only two copies of each gene are present.

### Bistability analysis and model calibration

To determine if our model exhibits bistability, we used the chemical reaction network Toolbox (23), which provides a set of kinetic constants that guarantee bistability along with the corresponding steady states. We then calibrated the full model by fixing the kinetic parameters as determined in (36), Table S1. We utilized COPASI (37), a software tool for simulating and analyzing biochemical networks, for both calibrations: for the stationary states (Fig. 2A) and for recovery time after treatment with the NF-κB inhibitor DHMEQ (Fig. 2B). To solve the model equations, we used the following parameters in COPASI: Relative Tolerance 1.0e^-6^, Absolute Tolerance 1.0e^-12^, and Maximum Internal Steps 1e^5^. The method used was ‘LSDODA’.

### Model simulation and landscape surface

All simulations were performed using COPASI (37). For the deterministic simulations, we employed the same methods and parameters as previously mentioned. For the stochastic simulations, we utilized tau-leaping, a variant of the Gillespie algorithm, setting epsilon (the interval during which multiple reactions can take place) to 1.0e^-3^ and the maximum internal steps to 100 million (1.0e^8^). To construct the epigenetic attractor landscape, we modified the MATLAB code from reference (38). In our adaptation, rather than randomly exploring positions in phase space, we performed one million 30-minute stochastic simulations using COPASI (37). These simulations began from positions evenly distributed across the intervals (0, 1.8e^5^) for p50 and (0, 6.0e^5^) for p65. These simulations were then imported to the MATLAB code.

### Validating Spearman correlation

We validated the correlations between p50 and p65 RNAs from our simulations using a cohort of 34 basal and 17 HER2^+^ untreated patients (25, 26). To ensure comparability, we randomly selected 34 basal and 17 HER2^+^ single-cell stochastic simulations from our set of 120. From each simulation, we randomly chose one pair of p50 and p65 RNA molecule numbers and calculated the correlation across the entire set. These pairs represent RNA levels from different cells at various moments. Figs. 3A and 3B compare these correlations with those from the cohort, while the numerical values are presented in Fig. 3C. Each time this procedure is repeated, a different correlation may be found. To avoid bias, we repeated the strategy for measuring correlations 100 times, including only sets where the correlation p-value was less than 5.0e^-3^.

## Supporting information

Supplementary files containing Figs. S1 to S3, Table S1, Legend for movies S1 and S2 and SI References.

Spontaneous transition from HER2+ to TNBC subtypes.

The unstable intermediate state provides a fluctuation-susceptible slow route along the weak eigenvectors.

## ACKNOWLEDGMENTS

We thank Alexander Hoffmann and Mary Sehl for their inspiring discussions about the paper results and perspectives, and Mariana Lopes for her assistance with figure design. Funding: FJPL was supported by CNPq - Brazilian Council for Scientific and Technological Development (200176/2022-6).

