## Supplementary files containing Figs. S1 to S3, Table S1, Legend for movies S1 and S2 and SI References. for "NF-κB Epigenetic Attractor Landscape Drives Breast Cancer Heterogeneity"

1 Developmental Biology and Dynamical Systems Group. Universidade Federal do Rio de Janeiro, Campus Duque de Caxias Professor Geraldo Cidade, Brasil.
2 Institute for Quantitative and Computational Biosciences (QCBio), University of California Los Angeles – UCLA.
3 Departamento de Biofísica e Biometria (DBB), Universidade Estadual do Rio de Janeiro (UERJ), Brasil.
4 Stem Cell Laboratory, Divisão de Laboratórios Especializados, Instituto Nacional de Câncer, Brazil

Corresponding author: *Francisco Lopes

**This PDF file includes:**

- Figs. S1 to S3
- Table S1
- Legend for movies S1 and S2.
- SI References

**Model Reactions**

| | Description | Reaction | Forward constants | Forward constants units | Backward constants | Backward constants units |
|---|---|---|---|---|---|---|
| 1 | NFkB dimer formation | p50 + p65 ↔ NFkB | 2.66319E-06 | pl^2/(min*#) | 1.69683E+00 | pl/min |
| 2 | NFkB binds p65 promoter | NFkB + N0p65 ↔ N1p65 | 8.90887E-04 | pl^2/(min*#) | 1.04883E+02 | pl/min |
| 3 | p65 RNA synthesis | N1p65 → N1p65 + RNAp65 | 5.62307E+01 | pl/min | | |
| 4 | p65 synthesis | RNAp65 → RNAp65 + p65 | 4.495E+01 | pl/min | | |
| 5 | NFkB binds p50 promoter | NFkB + N0p50 ↔ N1p50 | 9.02865E-01 | pl^2/(min*#) | 8.2909E+04 | pl/min |
| 6 | p50 RNA synthesis | N1p50 → N1p50 + RNAp50 | 3.70696E+01 | pl/min | | |
| 7 | p50 synthesis | RNAp50 → RNAp50 + p50 | 9.38E+00 | pl/min | | |
| 8 | NFkB binds SIP1 promoter | NFkB + N0SIP1 ↔ N1SIP1 | 8.90887E-04 | pl^2/(min*#) | 2.5257E+00 | pl/min |
| 9 | SIP1 RNA synthesis | N1SIP1 → N1SIP1 + RNASIP1 | 7.30761E+02 | pl/min | | |
| 10 | SIP1 synthesis | RNASIP1 → RNASIP1 + SIP1 | 1.1684E+02 | pl/min | | |
| 11 | NFkB binds SLUG promoter | NFkB + N0SLUG ↔ N1SLUG | 8.90887E-04 | pl^2/(min*#) | 7.925E+01 | pl/min |
| 12 | SLUG RNA synthesis | N1SLUG → N1SLUG + RNASLUG | 2.62313E+03 | pl/min | | |
| 13 | SLUG synthesis | RNASLUG → RNASLUG + SLUG | 1.1684E+02 | pl/min | | |
| 14 | NFkB binds TWIST promoter | NFkB + N0TWIST1 ↔ N1TWIST1 | 8.90887E-04 | pl^2/(min*#) | 4.85495E+00 | pl/min |
| 15 | TWIST RNA synthesis | N1TWIST1 → N1TWIST1 + RNATWIST1 | 8.08799E+03 | pl/min | | |
| 16 | TWIST synthesis | RNATWIST1 → RNATWIST1 + TWIST1 | 1.1684E+02 | pl/min | | |
| 17 | p50 RNA constitutive syntesis | 0 → RNAp50 | 6.12305E+00 | #/min | | |
| 18 | p65 RNA constitutive syntesis | 0 → RNAp65 | 9.19739E+00 | #/min | | |
| 19 | p50 RNA degradation | RNAp50 → 0 | 1.015E-01 | pl/min | | |
| 20 | p65 RNA degradation | RNAp65 → 0 | 1.07848E-01 | pl/min | | |
| 21 | SIP1 RNA degradation | RNASIP1 → 0 | 1.01208E-01 | pl/min | | |
| 22 | SLUG RNA degradation | RNASLUG → 0 | 1.01208E-01 | pl/min | | |
| 23 | TWIST1 RNA degradation | RNATWIST1 → 0 | 1.01208E-01 | pl/min | | |
| 24 | p65 degradation | p65 → 0 | 5.2506E-02 | pl/min | | |
| 25 | p50 degradation | p50 → 0 | 3.01325E-02 | pl/min | | |
| 26 | SIP1 degradation | SIP1 → 0 | 4.83112E-02 | pl/min | | |
| 27 | SLUG degradation | SLUG → 0 | 4.83112E-02 | pl/min | | |
| 28 | TWIST1 degradation | TWIST1 → 0 | 4.83112E-02 | pl/min | | |

**Fig. S1. Gene Regulatory Reactions and Parameters.** Blue parameters were calculated based on experimentally determined mRNA and protein half-lives, and transcription and translation rate constants (1). Green values were derived from the average corresponding values for a mammalian cell.

$$
\begin{aligned}
\frac{d([\mathrm{NFkB}])}{dt} = &+(k_{1_{\text{"NFkB dimer formation"}}} \cdot [p50] \cdot [p65] \\
&-k_{2_{\text{"NFkB dimer formation"}}} \cdot [\mathrm{NFkB}]) \\
&-(k_{1_{\text{"NFkB binds p50 gene promoter"}}} \cdot [\mathrm{NFkB}] \cdot [\mathrm{NO}p50] \\
&-k_{2_{\text{"NFkB binds p50 gene promoter"}}} \cdot [\mathrm{N1}p50]) \\
&-(k_{1_{\text{"NFkB binds p65 gene promoter"}}} \cdot [\mathrm{NFkB}] \cdot [\mathrm{NO}p65] \\
&-k_{2_{\text{"NFkB binds p65 gene promoter"}}} \cdot [\mathrm{N1}p65]) \\
&-(k_{1_{\text{"NFkB binds SIP1 gene promoter"}}} \cdot [\mathrm{NFkB}] \cdot [\mathrm{NO}SIP1] \\
&-k_{2_{\text{"NFkB binds SIP1 gene promoter"}}} \cdot [\mathrm{N1}SIP1]) \\
&-(k_{1_{\text{"NFkB binds SLUG gene promoter"}}} \cdot [\mathrm{NFkB}] \cdot [\mathrm{NO}SLUG] \\
&-k_{2_{\text{"NFkB binds SLUG gene promoter"}}} \cdot [\mathrm{N1}SLUG]) \\
&-(k_{1_{\text{"NFkB binds TWIST gene promoter"}}} \cdot [\mathrm{NFkB}] \cdot [\mathrm{NO}TWIST1] \\
&-k_{2_{\text{"NFkB binds TWIST gene promoter"}}} \cdot [\mathrm{N1}TWIST1])
\end{aligned} \tag{1}
$$

$$
\begin{aligned}
\frac{d([\mathrm{SLUG}])}{dt} = &-k_{1_{\text{"SLUG degradation"}}} \cdot [\mathrm{SLUG}] \\
&+k_{1_{\text{"SLUG synthesis"}}} \cdot [\mathrm{RNASLUG}]
\end{aligned} \tag{2}
$$

$$
\begin{aligned}
\frac{d([\mathrm{TWIST1}])}{dt} = &+k_{1_{\text{"TWIST synthesis"}}} \cdot [\mathrm{RNATWIST1}] \\
&-k_{1_{\text{"TWIST1 degradation"}}} \cdot [\mathrm{TWIST1}]
\end{aligned} \tag{3}
$$

$$
\begin{aligned}
\frac{d([\mathrm{RNATWIST1}])}{dt} = &-k_{1_{\text{"TWIST1 RNA degradation"}}} \cdot [\mathrm{RNATWIST1}] \\
&+ k_{1_{\text{"TWIST RNA synthesis"}}} \cdot [\mathrm{N1TWIST1}]
\end{aligned} \tag{4}
$$

$$
\begin{aligned}
\frac{d([\mathrm{RNASIP1}])}{dt} = &-k_{1_{\text{"SIP1 RNA degradation"}}} \cdot [\mathrm{RNASIP1}] \\
&+k_{1_{\text{"SIP1 RNA synthesis"}}} \cdot [\mathrm{N1SIP1}]
\end{aligned} \tag{5}
$$

$$
\begin{aligned}
\frac{d([\mathrm{RNASLUG}])}{dt} = &-k_{1_{\text{"SLUG RNA degradation"}}} \cdot [\mathrm{RNASLUG}] \\
&+k_{1_{\text{"SLUG RNA synthesis"}}} \cdot [\mathrm{N1SLUG}]
\end{aligned} \tag{6}
$$

$$
\begin{aligned}
\frac{d([\mathrm{p65}])}{dt} = &-k_{1_{\text{"p65 degradation"}}} \cdot [\mathrm{p65}] \\
&- k_{1_{\text{"NFkB dimer formation"}}} \cdot [p50] \cdot [p65] \\
&+k_{2_{\text{"NFkB dimer formation"}}} \cdot [\mathrm{NFkB}] \\
&+k_{1_{\text{"p65 synthesis"}}} \cdot [\mathrm{RNAp65}]
\end{aligned} \tag{7}
$$

**Fig. S2. Model differential equations.** System of Ordinary Differential Equations obtained by applying the Law of Mass Action to the model reactions in Fig. S1. Continue…

$$\frac{d([\text{RNAp65}])}{dt} = +k_{1_{\text{"p65 RNA synthesis"}}} \cdot [\text{N1p65}] - k_{1_{\text{"p65 RNA degradation"}}} \cdot [\text{RNAp65}] + \left(k_{1_{\text{"p65 RNA constitutive synthesis"}}}\right) \quad (8)$$

$$\frac{d([\text{RNAp50}])}{dt} = +k_{1_{\text{"p50 RNA synthesis"}}} \cdot [\text{N1p50}] - k_{1_{\text{"p50 RNA degradation"}}} \cdot [\text{RNAp50}] + \left(k_{1_{\text{"p50 RNA constitutive synthesis"}}}\right) \quad (9)$$

$$\frac{d([\text{p50}])}{dt} = -(k_{1_{\text{"NFkB dimer formation"}}} \cdot [p50] \cdot [p65] - k_{2_{\text{"NFkB dimer formation"}}} \cdot [\text{NFkB}]) - k_{1_{\text{"p50 degradation"}}} \cdot [p50] + k_{1_{\text{"p50 synthesis"}}} \cdot [\text{RNAp50}] \quad (10)$$

$$\frac{d([\text{NOSLUG}])}{dt} = -\left(k_{1_{\text{"NFkB binds SLUG gene promoter"}}} \cdot [\text{NFkB}] \cdot [\text{NOSLUG}] - k_{2_{\text{"NFkB binds SLUG gene promoter"}}} \cdot [\text{N1SLUG}]\right) \quad (11)$$

$$\frac{d([\text{NOSIP1}])}{dt} = -\left(k_{1_{\text{"NFkB binds SIP1 gene promoter"}}} \cdot [\text{NFkB}] \cdot [\text{NOSIP1}] - k_{2_{\text{"NFkB binds SIP1 gene promoter"}}} \cdot [\text{N1SIP1}]\right) \quad (12)$$

$$\frac{d([\text{SIP1}])}{dt} = -k_{1_{\text{"SIP1 degradation"}}} \cdot [\text{SIP1}] + k_{1_{\text{"SIP1 synthesis"}}} \cdot [\text{RNASIP1}] \quad (13)$$

$$\frac{d([\text{NOTWIST1}])}{dt} = -\left(k_{1_{\text{"NFkB binds TWIST gene promoter"}}} \cdot [\text{NFkB}] \cdot [\text{NOTWIST1}] - k_{2_{\text{"NFkB binds TWIST gene promoter"}}} \cdot [\text{N1TWIST1}]\right) \quad (14)$$

$$\frac{d([\text{N1TWIST1}])}{dt} = +k_{1_{\text{"NFkB binds TWIST gene promoter"}}} \cdot [\text{NFkB}] \cdot [\text{NOTWIST1}] - k_{2_{\text{"NFkB binds TWIST gene promoter"}}} \cdot [\text{N1TWIST1}] \quad (15)$$

$$\frac{d([\text{N1SLUG}])}{dt} = +k_{1_{\text{"NFkB binds SLUG gene promoter"}}} \cdot [\text{NFkB}] \cdot [\text{NOSLUG}] - k_{2_{\text{"NFkB binds SLUG gene promoter"}}} \cdot [\text{N1SLUG}] \quad (16)$$

$$\frac{d([\text{N1SIP1}])}{dt} = +k_{1_{\text{"NFkB binds SIP1 gene promoter"}}} \cdot [\text{NFkB}] \cdot [\text{NOSIP1}] - k_{2_{\text{"NFkB binds SIP1 gene promoter"}}} \cdot [\text{N1SIP1}] \quad (17)$$

$$\frac{d([\text{N1p65}])}{dt} = +k_{1_{\text{"NFkB binds p65 gene promoter"}}} \cdot [\text{NFkB}] \cdot [\text{NOp65}] - k_{2_{\text{"NFkB binds p65 gene promoter"}}} \cdot [\text{N1p65}] \quad (18)$$

$$\frac{d([\text{N1p50}])}{dt} = +k_{1_{\text{"NFkB binds p50 gene promoter"}}} \cdot [\text{NFkB}] \cdot [\text{NOp50}] - k_{2_{\text{"NFkB binds p50 gene promoter"}}} \cdot [\text{N1p50}] \quad (19)$$

$$\frac{d([\text{NOp65}])}{dt} = -\left(k_{1_{\text{"NFkB binds p65 gene promoter"}}} \cdot [\text{NFkB}] \cdot [\text{NOp65}] - k_{2_{\text{"NFkB binds p65 gene promoter"}}} \cdot [\text{N1p65}]\right) \quad (20)$$

$$\frac{d([\text{NOp50}])}{dt} = -\left(k_{1_{\text{"NFkB binds p50 gene promoter"}}} \cdot [\text{NFkB}] \cdot [\text{NOp50}] - k_{2_{\text{"NFkB binds p50 gene promoter"}}} \cdot [\text{N1p50}]\right) \quad (21)$$

**Fig. S2. Model differential equations.** System of Ordinary Differential Equations obtained by applying the Law of Mass Action to the model reactions in Fig. S1. *End.*

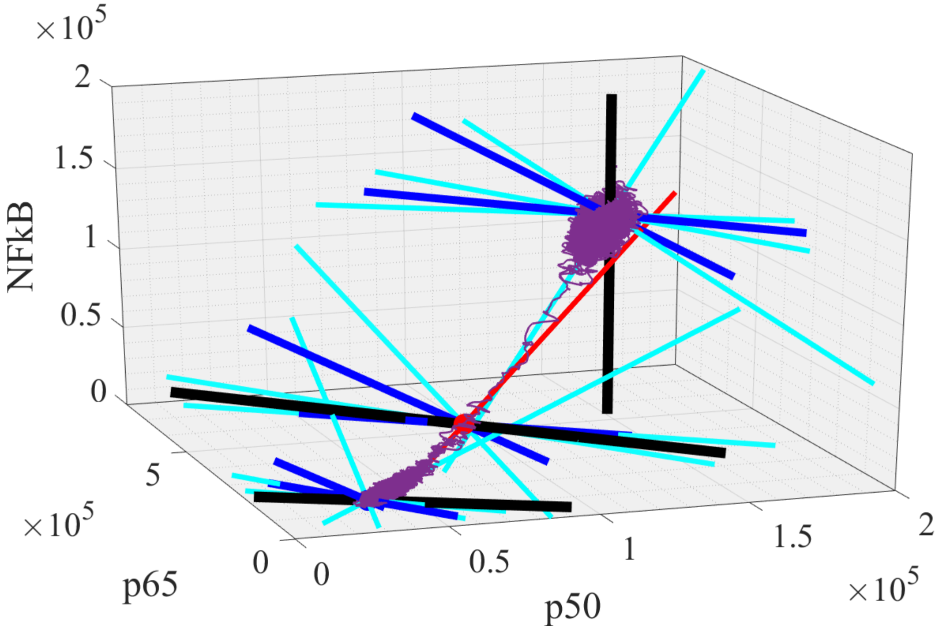


**Fig. S3. Eigenvectors.** Projection of all seven eigenvectors with nonzero eigenvalues, determined from each stationary state, within the p65-p50-NFκB phase-space. The magnitude of the associated eigenvalues varies widely, ranging from –2.0e5 to 2.5e-3. To deal with this wide range, we classified the eigenvectors into three categories based on their eigenvalues: light blue for weak ($1.0e^{-3}$ to $1.0e^{-1}$), dark blue for medium (1.0 to 1.0e+2), and black for strong ($1.0e^{+4}$ or higher). The only eigenvector with a positive eigenvalue is the red one, which is classified as weak ($2.5e^{-3}$). The trajectory from a single-cell stochastic simulation is shown in purple.

**Table S1. mRNA and protein copy number estimates**. mRNA copy number was estimated by multiplying the number of molecules in a healthy cell (1) by the relative increase given by the qPCR data. Protein copy number was estimated from the RNA copy number, the mRNA translation rate constant, and the protein degradation rate (1). TWIST1, SLUG, and SIP1 mRNA molecules were estimated from the median of all values in a healthy cell. The same approach was used for their mRNA translation rate constant and protein degradation rate. NFκB1 (p50) was estimated from the RelA (p65) qPCR data.

| | HER2 cell | | | | TNBC cell | | | |
|---|---|---|---|---|---|---|---|---|
| | qPCR | | mRNA copy number | Protein copy number | qPCR | | mRNA copy number | Protein copy number |
| | Mean | Standard Error | | | Mean | Standard Error | | |
| RelA | 3.33 | $3.94e^{-1}$ | $1.35e^{2}$ | $1.16e^{5}$ | $1.54e^{1}$ | $6.84e^{1}$ | $6.25e^{2}$ | $5.35e^{5}$ |
| NF-κB1 | | | $1.04e^{2}$ | $3.25e^{4}$ | | | $4.83e^{2}$ | $1.50e^{5}$ |
| TWIST1 | $4.78e^{3}$ | $6.26e^{2}$ | $8.30e^{4}$ | $2.01e^{8}$ | $8.81e^{3}$ | $2.78e^{2}$ | $1.53e^{5}$ | $3.71e^{8}$ |
| SLUG | $1.85e^{2}$ | $5.30e^{1}$ | $3.22e^{3}$ | $7.79e^{6}$ | $1.75e^{3}$ | $4.35e^{1}$ | $3.04e^{4}$ | $7.36e^{7}$ |
| SIP1 | $5.61e^{2}$ | 4.04 | $9.75e^{3}$ | $2.36e^{7}$ | $8.12e^{2}$ | $1.63e^{2}$ | $1.41e^{4}$ | $3.42e^{7}$ |

**Movie S1 (separate file). Spontaneous transition from HER2+ to TNBC subtypes.** The purple line indicates the full trajectory in the p50-p65 phase space. The time-lapse animation shows fluctuations within each attractor basin as well as the spontaneous transition.

**Movie S2 (separate file). The unstable intermediate state provides a fluctuation-susceptible slow route along the weak eigenvectors.** The time-lapse animation shows that the protein copy number fluctuate along the weak eigenvectors in the p50-p65 phase space. The full trajectory is depicted in Figure 4A.
